# FITNESS BENEFITS TO BACTERIA OF CARRYING PROPHAGES AND PROPHAGE-ENCODED ANTIBIOTIC-RESISTANCE GENES PEAK IN DIFFERENT ENVIRONMENTS

**DOI:** 10.1101/2020.03.13.990044

**Authors:** Carolin C. Wendling, Dominik Refardt, Alex R. Hall

## Abstract

Bacteria can acquire antibiotic resistance genes (ARGs) via prophages, phage genomes integrated into bacterial chromosomes. Such prophages may influence bacterial fitness via increased antibiotic resistance, protection from further phage infection, or by switching to a lytic lifecycle that releases free phages which can infect phage-susceptible competitors. We expect these effects to depend on environmental conditions because of, for example, environment-dependent induction of the lytic lifecycle. However, our understanding of how costs and benefits of prophage-encoded ARGs vary across environments remains limited. Here, by studying prophages with and without ARGs in *Escherichia coli*, we distinguished between effects of prophages alone and ARGs they carry. In competition with prophage-free strains, fitness benefits from prophages and ARGs peaked in different environments. Prophage carriage was most beneficial in conditions where induction of the lytic lifecycle was common, whereas ARGs were more beneficial in the presence of antibiotics and when prophage induction was lower. Acquisition of prophage-encoded ARGs by competing phage-susceptible strains was most common when prophage induction, and therefore the amount of free phages, was high. Thus, selection on prophages and ARGs varies independently across environments, which is important for predicting the spread of mobile/integrating genetic elements and their role in antibiotic resistance evolution.

## Introduction

Evolution of antimicrobial resistance (AMR) is a natural process that occurs due to *de novo* mutation or horizontal gene transfer (HGT). HGT of antibiotic resistance genes (ARGs) can occur within and across species by conjugation, transformation or transduction (transfer of foreign DNA by independently replicating bacteriophages, phages). By facilitating the exchange of genetic material through HGT, phages are widely recognized as playing a significant role in bacterial adaptation in general (Rohwer 2003; Mann 2005) and may be important during bacterial adaptation to antibiotics, potentially acting as “vehicles for resistance genes” (Balcazar 2014). In support, recent high-throughput sequencing suggested phages represent environmental reservoirs of ARGs (Calero-Caceres et al. 2019; Debroas and Siguret 2019). However, the contribution of phages, and in particular prophages (phages integrated into bacterial chromosomes) to the acquisition, maintenance and spread of ARGs remains incompletely understood. Despite evidence that prophages can encode ARGs (Brenciani et al. 2010; Palmieri et al. 2011), it is not clear how carriage of such prophages affects bacterial fitness, how these effects depend on the presence of the ARGs, and how they vary with local environmental conditions. Quantifying the net fitness effect of prophages with and without ARGs on their bacterial hosts in different environments will therefore improve our basic understanding of bacteria-virus interactions and the potential relevance of prophage-mediated antibiotic resistance evolution for infectious diseases in humans and animals.

We hypothesize the net fitness effect of an ARG-carrying prophage for its bacterial host varies strongly depending on environmental conditions. Prophages are formed when temperate phages incorporate their viral genome into the bacterial chromosome (unlike lytic phages, which multiply inside the host cell before lysing it and releasing new virions). The prophage then replicates as part of the bacterial host cell (now called a lysogen). In this state, prophages can confer benefits to their hosts, for example by conferring immunity against superinfecting phages (Refardt 2011) or providing them with beneficial genes, including ARGs (Modi et al. 2013). We expect the benefit of carrying a prophage-encoded ARG relative to costs of expressing it will be higher in the presence of the selecting antibiotic, whereas in the absence of the antibiotic, expression of the ARG may have a net negative effect on bacterial fitness (a ‘cost of resistance’ (Andersson and Levin 1999)). In some conditions, we expect such costs to be compounded by prophages switching back to the lytic cycle and killing the host (Casjens 2003; Paul 2008). This switch from the lysogenic to the lytic cycle can happen spontaneously or in response to environmental conditions, such as sub-inhibitory antibiotic concentrations that trigger the bacterial SOS response (Otsuji et al. 1959). Although lysis is detrimental for individual cells, the prophage-carrying population might in fact benefit from the release of free phages, because they may infect and kill any competing phage-susceptible cells in the same microbial community (Ubeda et al. 2005; Andersson and Hughes 2014; Haaber et al. 2016). The ultimate impact of prophages on their host’s fitness is complicated further by the possibility that any free phages released via lysis can potentially transfer ARGs to competing phage susceptible cells, further modulating competition between bacterial populations. Consequently, we expect the fitness of lysogens relative to prophage-free strains to vary depending on antibiotic concentration and other environmental factors influencing prophage induction and lysogeny.

Fitness costs and benefits associated with prophages that encode ARGs remain incompletely understood. This might be due, among other reasons, to the difficulty of directly comparing prophage-carrying with prophage-free bacteria, thereby accounting for the presence/absence of ARGs encoded on prophages. Consequently, the following questions remain unanswered: (1) How does the net fitness effect of prophage carriage vary across different antibiotic concentrations? (2) Are these effects specific to prophages that encode ARGs? (3) Are prophages more beneficial in environments where lysis is more frequent? (4) In which environments is the acquisition of ARG-carrying prophages by susceptible strains most likely? To answer these questions, we determined fitness costs and benefits associated with three common ARGs encoded on four different naturally occurring prophages of *Escherichia coli*. We used strains carrying these prophages and ARGs in various combinations (12 different lysogens), including those where the prophage encoded an ARG and where it did not, and competed them against an antibiotic- and phage-susceptible strain. We performed competitions (more than 600 in total) in multiple environments (presence and absence of selective antibiotics at various concentrations and of compounds that induce the lytic cycle of the prophage). This allowed us to differentiate between the effects of prophages alone and prophages carrying ARGs on their bacterial host’s fitness, and to do so across antibiotic concentrations and conditions associated with increased lysis and free phage production.

## Material and Methods

### Organisms

We used *E. coli* K-12 MG1655 (hereafter Wild Type, WT) and 12 different lysogens constructed from this strain (Table 1). Specifically, we used four different prophages, lambda+ (originally obtained as strain JL573 from J.W. Little, University of Arizona), HK022, Phi80 (both originally obtained from M. E. Gottesman, Columbia University) and mEp234 (originally obtained from I.-N. Wang, SUNY Albany). The prophage in each lysogen encoded either no antibiotic resistance gene (ARG) or one of three different ARGs conferring resistance to ampicillin (amp), kanamycin (kan) or chloramphenicol (cm), all of which have been widely detected among naturally occurring coliphages (Shousha et al. 2015).

**Table 1.**
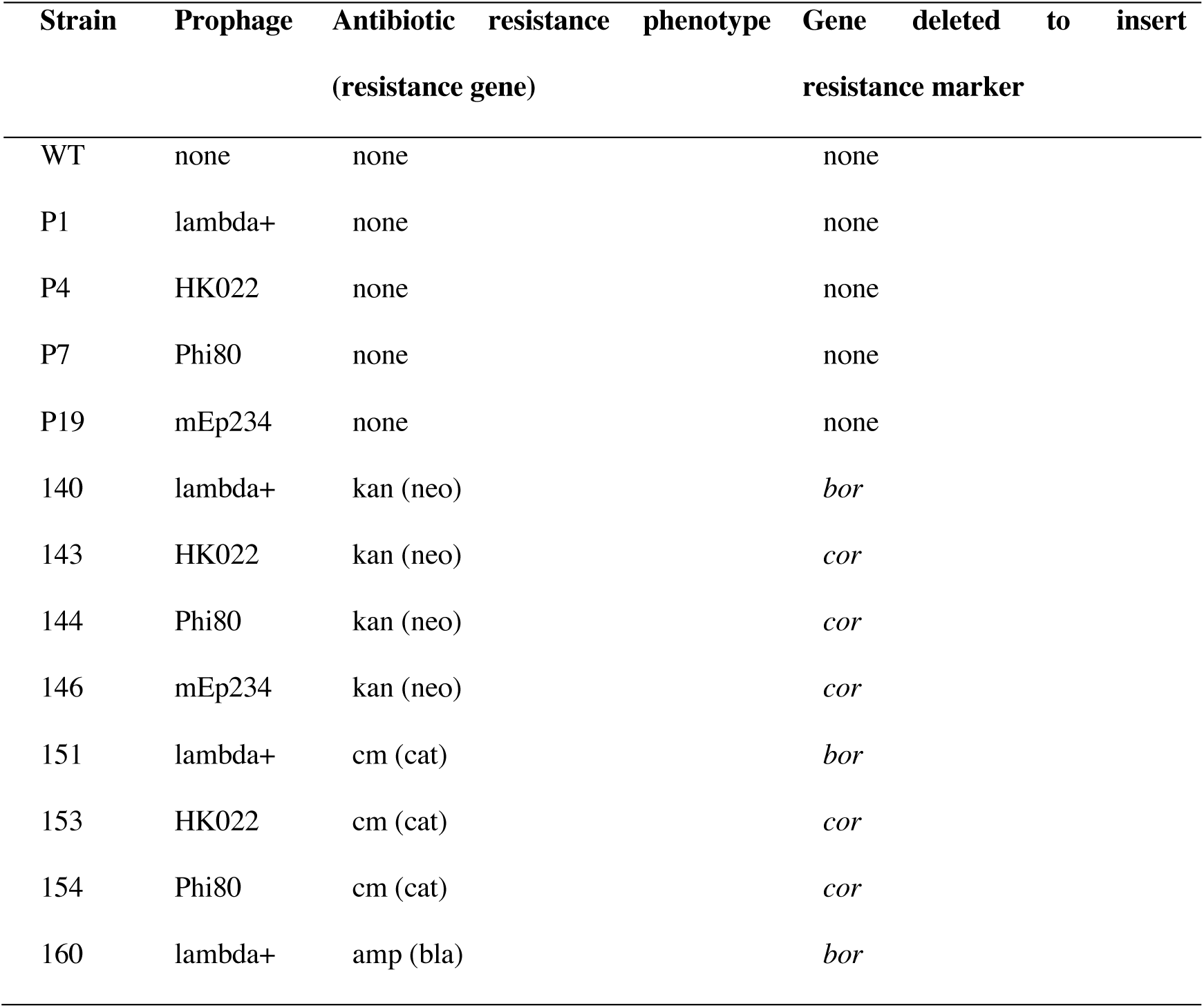
Bacterial strains. The Wild Type (WT) is *E. coli* K-12 MG1655. All other strains are lysogens constructed from this strain. Antibiotic resistance phenotype abbreviations: amp - ampicillin, cm - Chloramphenicol, kan - kanamycin. Resistance gene abbreviations: *bla* - beta-lactamase, *cat* - chloramphenicol acetyltransferase, *neo* - aminoglycoside phosphotransferase from Tn5.

Phages carrying an ARG were constructed using the first step (exchange of a gene with a resistance cassette) of a λ Red-promoted gene replacement (Datsenko and Wanner 2000)0). Resistance cassettes were amplified from plasmids pKD3 (*cat*, chloramphenicol resistance), pKD4 (*neo*, kanamycin resistance), and pGEM®-T Easy (*bla*, ampicillin resistance) with primers that carry 5’ homology extensions (Supplementary material, Table S1). Primers were designed to omit the FRT-sites on pKD3 and pKD4. Lysogens were then transformed with PCR products by electroporation. To enable a recombination event with the prophage, lysogens carried the plasmid pKD46, a temperature-sensitive low-copy number plasmid that expresses the λ Red genes under the control of an arabinose-inducible promoter and were induced with 20 mM arabinose prior to electroporation. Transformed cells were subsequently plated on the respective antibiotic and successful transformants were isolated, induced and the recombineered phages were inserted into fresh WT cells.

### Determination of optimal antibiotic concentrations to use in competition experiments

We measured bacterial growth of each strain across an antibiotic gradient to identify suitable antibiotic concentrations for the competition assays and to determine whether prophage carriage alone (without ARGs) had an effect on antibiotic resistance. Growth assays were performed for each strain at 37 °C in triplicates in 96-well plates containing ten different antibiotic concentrations, no-drug controls and contamination controls (medium only). At the beginning (t=0) and after 20 hours (t=20) we measured optical density at 600 nm (OD600). We fitted a Hill-function to the data (Regoes et al. 2004) of the change in optical density (OD_t=20_ – OD_t=0_) as a function of antibiotic concentration and extracted different inhibitory concentrations (ICs), at which growth relative to that observed for the same strain in the absence of the antibiotic was reduced by 20% (IC20), 30% (IC30) and 50% (IC50). We confirmed that the ARGs encoded on prophages conferred antibiotic resistance to lysogens by observing much higher ICs for ARG-carrying lysogens compared to the WT (Figure S1). In preliminary competition assays, we observed extinction of the WT at concentrations higher than IC50. We therefore performed the main competition experiment at the following antibiotic concentrations: high (IC50 for the WT; Kan: 2.17*µ*g/ml, Cm: 1.31 *µ*g/ml, Amp: 4.41 *µ*g/ml), intermediate (IC30 for the WT; Kan: 1.55 *µ*g/ml, Cm: 1.08 *µ*g/ml, Amp: 3.52 *µ*g/ml) and low (IC20 for the WT; Kan: 0.565 *µ*g/m, Cm: 0.623 *µ*g/ml, Amp: 1.8*µ*g/ml).

### Competition Assays

To determine the costs and benefits for bacteria of carrying prophages we performed competition assays between each lysogen (Table 1) and a phenotypically marked version of the Wild Type. This competing strain is susceptible to each phage and antibiotic, and is labelled with a fluorescent yellow-super-fluorescent protein (SYFP) marker (Gullberg et al. 2014). For the eight lysogens where the prophage encoded an ARG (Table 1), we assayed each lysogen with and without the respective antibiotic at each concentration. For each ARG-carrying lysogen, we also assayed a corresponding ARG-free lysogen (Table 1; carrying the same prophage but without an ARG) in the same conditions. We tested each of these combinations of lysogen (*n*=8), ARG (with/without) and antibiotic concentration (zero, low, intermediate, high) in the presence and absence of a compound that induces the lytic cycle (mitomycin C, at a final concentration of 0.5 *µ*g/ml). With five replicates, this gave 640 competitions in total. We also accounted for the effect of the SYFP marker carried by the susceptible competing strain, by measuring the fitness of the SYFP-labelled WT relative to the unlabelled WT in each of the tested environments. We then subtracted the difference in relative fitness between both WT strains from the fitness of each lysogen relative to the YFP-labelled WT for each environment.

For each competition assay, single colonies of each competing clone were inoculated in 5 ml of LB and grown overnight at 37 °C in a shaking incubator (180 rpm). The following day we determined the optical density of each overnight culture at 600 nm and adjusted the OD of each culture to the lowest measured OD, to obtain similar concentrations of bacteria per culture. Then, cultures of each lysogen were mixed 1:1 (*v:v*) with the SYFP-labelled WT, diluted 1:1000 into the respective environment (combination of antibiotic concentration and mitomycin C) in a 96-well microplate to initiate each competition culture, and incubated overnight at 37 °C. At the start of each competition and after 24 h, absolute cell densities of the two competing populations were determined by flow cytometry (Novosampler Pro; ACEA Biosciences Inc., San Diego CA), where fluorescent (WT) and non-fluorescent cells (lysogens) were counted, similarly to past work on costs and benefits of antibiotic resistance (Qi et al. 2016). Competitive fitness was estimated as selection rate constant, *r*, calculated as the difference of the realized Malthusian parameters between competing strains (Lenski et al. 1991):

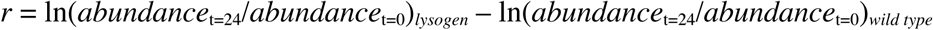

### Detection of prophage acquisition by the Wild Type

We plated the competing-cultures after 24 h onto non-selective and selective agar-plates (supplemented with the respective antibiotic) to determine the presence and frequency of newly antibiotic-resistant WT clones (which could grow on antibiotic plates and were distinguished from the competing strain by yellow fluorescence). We did this only for competition cultures involving ARG-carrying lysogens. To determine whether newly antibiotic-resistant colonies had acquired antibiotic resistance by lysogenization (acquiring the prophage), all antibiotic-resistant WT clones were subsequently checked for the presence of the respective prophage using prophage-induction by mitomycin C. This works by incubating 100 *µ*l of bacterial culture with 0.5 *µ*g/ml mitomycin C for 4 hours in a spectrophotometer, measuring optical density periodically. If prophages have been acquired, we expect the population to crash and eventually clear completely due to prophage induction and lysis, which in these conditions typically occurs after ∼1-2 hours (Refardt 2011). The supernatant of these mitomycin C-induced cultures was further spotted on a lawn of susceptible *E. coli* K12 MG1655, to confirm the presence of free phages in the induced cultures (see also below in section *Production of free phages*). We also used a second test for evidence of prophage acquisition, by assaying resistance of each putative new lysogen to a lytic version of the respective prophage (prophage acquisition is expected to confer immunity against superinfection by the same phage genotype). We did this by cross-streaking each putative new lysogen over a line of the respective lytic phage (Refardt 2011), and inferred resistance when new lysogens showed no visible sign of inhibition by phage after 24 h incubation.

We estimated relative fitness of the new lysogens (WT clones that acquired antibiotic resistance via lysogenization during the competition assay) by competition assays as above. We did this for each new lysogen against the ancestral WT (telling us about the net effect of acquiring the prophage including the ARG), the ancestral lysogen without the respective ARG (telling us about the effect of the ARG), and the ancestral lysogen with the ARG (which we would expect to have similar fitness to the new lysogens). Competitive fitness assays were performed in triplicate in the absence of antibiotics and at the IC20 of the respective antibiotic as above.

### Production of free phages

Lysogens can release free phages due to spontaneous prophage induction or in response to environmental stress, such as exposure to mitomycin C (Pricer and Weissbach 1964). In addition to the competition assays, we determined the production of free phages for each tested lysogen in the same eight environments used during competition assays (four antibiotic concentrations each tested in the presence and absence of mitomycin C) by means of standard spot-assays (Carlson 2005) in a separate experiment. Overnight cultures of each lysogen (with and without ARGs), grown from single colonies, were diluted 1/1000 in 5 ml LB and incubated for 2 h to bring cultures into exponential growth. Each of these 2 h-growth cultures was adjusted to a similar optical density (OD 600 nm) to achieve approximately equal cell densities, before 180 *µ*l was transferred to a 96-well microplate. After 4 h (when induction had completed, which was visible by complete clearance of mitomycin C-treated cultures), we diluted each culture 1/10 in SM-Buffer, added a drop of chloroform and centrifuged for 10 min at 5000 *g* to remove bacteria and isolate any free phages. To estimate the abundance of free phages in each culture, we then spotted a dilution series of these lysates on double-agar plates containing the WT in the overlay agar. Plates were incubated overnight at 37 °C and plaque forming units (PFU) were counted the following day, with three replicates per treatment.

### Reduction in bacterial growth caused by free phages

We measured the ability of free phages to inhibit population growth of the WT by determining the reduction in bacterial growth (RBG) in liquid culture, adapted from (Poullain et al. 2008). To do this we first isolated free phages from each lysogen (Table 1), by induction as described above using 0.5 *µ*g/ml mitomycin C in the absence of antibiotics. We then incubated cultures of the WT in 96-well plates, with each culture containing either no antibiotics or one of the three antibiotics (ampicillin, chloramphenicol or kanamycin) at IC20, IC30 or IC50. Each culture was inoculated at a concentration of ∼1.3×10^8^ CFU/ml of bacteria, to which we added ∼6.7×10^8^ PFU/ml of one of the phages or no phage (control). We then estimated bacterial abundance by measuring optical density at 600 nm at *t*=0 h and again at *t*=24 h of static incubation at 37 °C, with three replicates per treatment. The reduction in bacterial growth was calculated as:

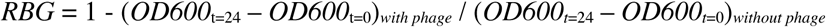

### Statistical analyses

All analyses were performed in R version 3.5.3.

#### Relative fitness

Our experimental design was not fully-factorial (our strains represent a subset of all possible phage×ARG combinations; Table 1). We therefore split the data set by antibiotic, which allowed us to fit a fully factorial linear model within each antibiotic treatment. In a first model, we took lysogen fitness relative to the WT in the absence of mitomycin C as the response variable, and as fixed effects we included antibiotic concentration (zero, IC20, IC30, IC50), ARG (with/without), and phage (Lambda, HK022, Phi80 or mEp234 for kanamycin; Lambda, HK022 or Phi for chloramphenicol; phage not included for ampicillin because only Lambda was represented with this antibiotic; Table 1). In a second model, we included the treatment groups with mitomycin C added, and mitomycin C (presence/ absence) as a fixed factor. In both models, we initially included all interaction terms, before reducing each model by sequentially removing non-significant interactions.

To separate the fitness effects of prophage carriage from the fitness effects of ARGs, we estimated the average fitness effect of each prophage in each treatment group (combination of antibiotic, antibiotic concentration, mitomycin C and prophage type) as the mean selection rate constant for the ARG-free lysogen relative to the WT. We then estimated the average fitness effect of each prophage-encoded ARG in each treatment group as the difference in mean selection rate constant between each ARG-carrying lysogen and the corresponding ARG-free lysogen in the same treatment group. That is, the fitness effect of the ARG here is the additional cost/benefit of the ARG after accounting for any fitness effect of the prophage it is carried on. Next, to determine which treatment groups were associated with similar fitness benefits of prophages and ARGs, we performed a pairwise-distance cluster analysis robust to outliers. We clustered treatment groups according to the variables (i) mean prophage fitness effect and (ii) mean ARG fitness effect. We did this separately for each antibiotic. Preliminary analysis of within-cluster sums of squares for different levels of *k* (the number of clusters) suggested that two to five clusters gave good levels of explanatory power for each antibiotic, so we proceeded with *k*-means clustering using km=3 in the R package *ComplexHeatmap*.

#### Production of free phages in each environment and absolute cell densities

Absolute cell densities during the pairwise competitions and free phage production of each lysogen in the same conditions were analysed following the same approach as for relative fitness data, using a log-transformation for free phage abundances (measured in PFU/ml). We further performed a correlation analysis to determine whether average relative fitness of lysogens in different treatment groups was correlated with the amount of free phages produced by lysogens in the same conditions (but measured in a separate experiment).

#### Phage predation rate

We determined differences in RBG for the different phages for each antibiotic using a linear model with antibiotic concentration, presence/absence of the respective ARG, phage type and all their interactions as fixed effects, reducing the model as above.

#### Lysogenization

Using data on the fraction of competition assays where we observed new lysogens, we used logistic regression to estimate the probability for the WT population to acquire an ARG-carrying prophage (lysogenization) in each environment (treatment group). We took presence/absence of new lysogens in each competition culture as the dependent variable and presence/absence of mitomycin C, antibiotic concentration (zero, IC20, IC30 or IC50), and phage type as independent variables. We pooled the data from the different antibiotic treatments here, because we found no effect of antibiotic (and therefore the type of resistance gene) on the probability to acquire an ARG-carrying prophage (glm: χ2(df=2) = 0.7, p=0.71). The full model was analysed by a generalized linear model and Analysis of Deviance, for which we assumed deviance change to be approximately χ2 distributed.

#### Relative fitness of WT-lysogens

To analyse the fitness of new lysogens (WT colony isolates that acquired an ARG-carrying prophage during the competition experiment), we used a linear mixed effects model (package: nlme, function lme) for each antibiotic (Amp, Cm, and Kan) with prophage type, presence/absence of antibiotics (IC20), and competitor (Phage-and-ARG-free WT, ARG-free lysogen, and ARG-carrying lysogen) and their interactions as fixed effects, and lysogen as random effect.

#### Growth rate in monoculture

We used the *SummarizeGrowth* function implemented in the R package *growthcurver* (Sprouffske and Wagner 2016) to estimate the intrinsic growth rate for each strain in the absence of antibiotics or mitomycin C (Table 1). We used Welch’s pairwise *t*-tests with sequential Bonferroni correction to determine whether individual lysogens had different growth rates compared to the WT.

## RESULTS

### Net benefits of ARG-encoding prophages increase at higher antibiotic concentrations

In the absence of antibiotics and mitomycin C, prophage-carrying strains (lysogens) had competitive fitness greater than or similar to the WT (selection rate constant ≥0; Figure 1a). In these conditions, there was no average difference between lysogens where the prophage encoded an ARG and those where it did not (paired *t*-test: t_7_=0.45, p=0.67; Figure 1a). This suggests neither ARGs nor prophages were costly in the absence of antibiotics. This was supported by experiments in monoculture, where we observed similar growth rates for lysogens compared to the WT (mean difference of lysogen growth rate from WT = 3.8% ±16.6% (s.d.); Figure S2). In the presence of antibiotics (but without mitomycin C), average lysogen fitness in our competition experiment increased (significant antibiotic concentration effect in linear model without mitomycin C: ampicillin treatment group: F_3,32_=104.255, p<0.001; chloramphenicol treatment group: F_3,96_=71.61, p<0.001; kanamycin treatment group: F_3,128_=18.04, p<0.001; Figure 1a). This was particularly pronounced for lysogens with ARG-carrying prophages, indicating resistance genes became advantageous at higher antibiotic concentrations (significant antibiotic concentration × ARG interaction in linear model for mitomycin C-free treatment groups: ampicillin: F_3,32_=18.03, p<0.001, chloramphenicol: F_3,96_=47.19, p<0.001, kanamycin: F_1,128_=13.56, p<0.001; Figure 1a). While this increase was qualitatively consistent across all ARG-encoding prophages, the fitness advantage of the ARG (relative to ARG-free versions of the same prophages) was much higher for chloramphenicol-resistance than for kanamycin- or ampicillin resistance (Figure 1a).

**Figure 1.**
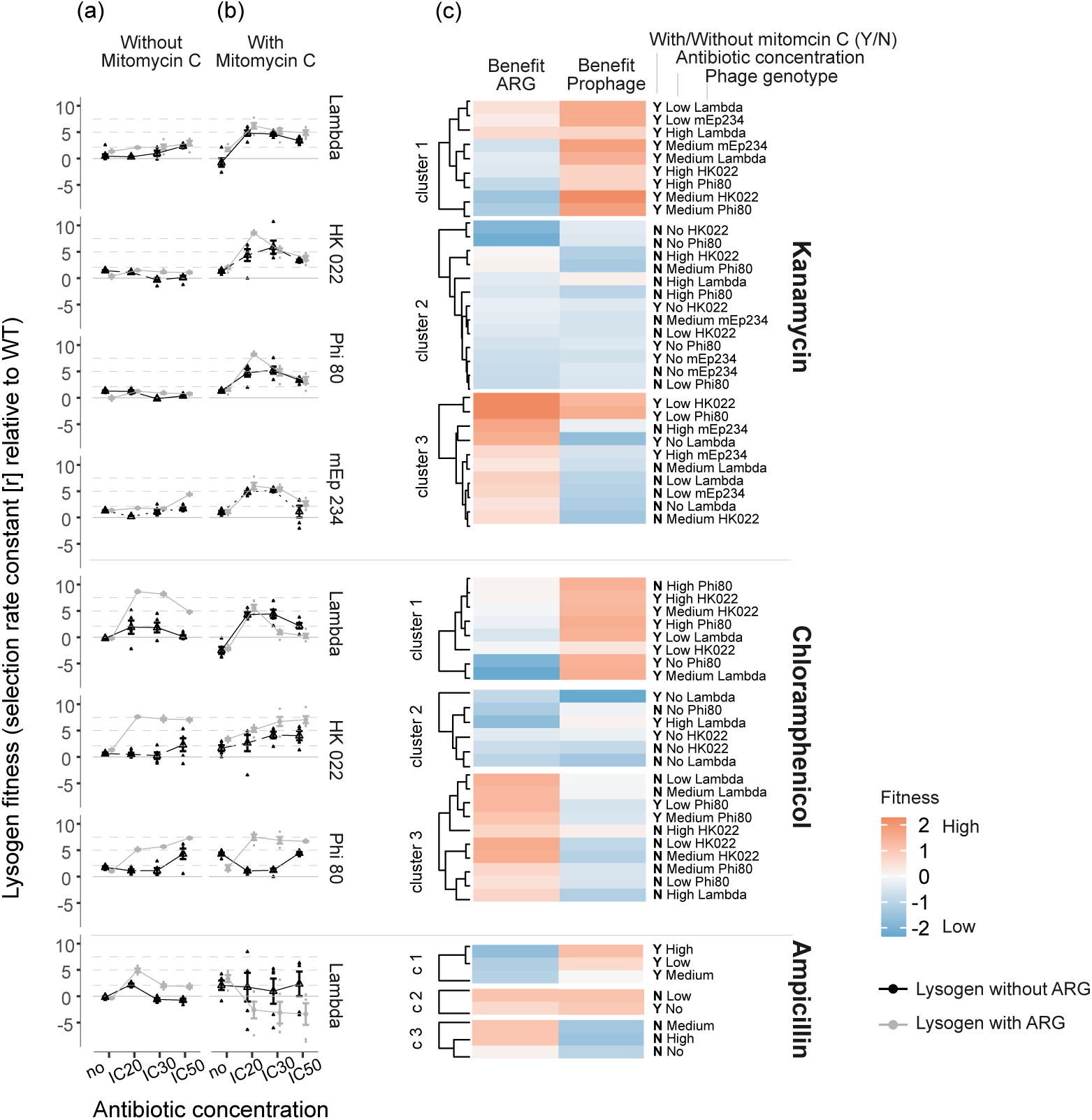
Competitive fitness of lysogens relative to the antibiotic-susceptible and phage-susceptible Wild Type in the (a) absence and (b) presence of mitomycin C. Each row of panels shows data from a different combination of antibiotic (given at right of panel (c)) and prophage type (given at right of panel (b)). Shown are single data points as well as means ± s.e. from six replicate populations. Black/grey points indicate lysogens without/with ARGs, assayed in competition with the WT at different antibiotic concentrations (*x*-axis). Lysogens have higher fitness than the WT when the selection rate constant *r* > 0. (c) Benefits of ARGs (left column) and prophages (right column) in different treatment groups (labelled at the right of the heatmap), clustered by *k*-means clustering (see Methods). Data have been scaled using the scale function implemented in R prior to the cluster analysis.

### Benefits of ARGs and prophages peak in different environments

In the presence of mitomycin C and at non-zero antibiotic concentrations, lysogens had on average higher competitive fitness relative to the WT than in the absence of mitomycin C (mitomycin C × antibiotic concentration interaction for ampicillin treatment group: F_3, 70_ = 4.21, p<0.001 and kanamycin treatment group: F_3, 245_ = 109.18, p<0.001; Figure 1b). This increase in lysogen fitness in the presence of mitomycin C and antibiotics was less pronounced for chloramphenicol (mitomycin C × antibiotic concentration interaction for chloramphenicol treatment group: F_3, 190_ = 0.55, p=0.64) and only significant for one intermediate chloramphenicol concentration (mitomycin C × antibiotic concentration=IC30 interaction: Estimate = 2.88, t=2.65, p=0.009). Overall, in the presence of mitomycin C, lysogen fitness peaked at intermediate antibiotic concentrations (IC20 or IC30) and declined with increasing antibiotic concentrations (with a few exceptions; Figure 1b).

The increasing benefits of ARGs at non-zero antibiotic concentrations observed above (Figure 1a) were modified by addition of mitomycin C (mitomycin C × ARG × antibiotic concentration interaction for chloramphenicol: F_3,190_ = 7.91, p<0.001 and kanamycin: F_3,254_ = 9.13, p<0.001; Figure 1b). This was because in the absence of mitomycin C and at non-zero antibiotic concentrations, carriage of an ARG had a larger effect on lysogen fitness than carrying a prophage did (in Cluster 3 there was a higher average benefit of ARGs than prophages, and these treatment groups were mostly mitomycin C-free; Figure 1c). In contrast, in the presence of mitomycin C, the fitness benefits of ARGs relative to prophages were less pronounced or even absent, and lysogen fitness was primarily determined by prophage carriage (in Cluster 1 there was a higher benefit of prophages than ARGs, and mitomycin C was present in most of these treatment groups; Figure 1c). Crucially, this pattern of relatively large benefits of prophage carriage in the presence of mitomycin C, and of ARG carriage in the absence of mitomycin C and presence of antibiotics, was consistent across all three antibiotics (Figure 1c). The reduced benefit of ARG carriage was particularly pronounced in the presence of ampicillin and mitomycin C, where Amp^R^-lysogens were at a disadvantage relative to the WT (Figure 1b-c).

We observed no significant variation of mean lysogen fitness among the four different prophage types (lambda, HK022, Phi80, mEp234) in the presence of kanamycin (phage-genotype effect for kanamycin: F_3, 250_ = 1.59, p=0.19). However, in the presence of chloramphenicol, average lysogen fitness varied among prophage types (prophage type effect for chloramphenicol: F_2,186_ = 37.33, p<0.001; Figure 1). This indicates the variation of lysogen fitness across different types of ARG-carrying prophages depends on which type of antibiotic resistance we look at.

### Increased lysogen fitness in environments that support higher free-phage production

When grown in monoculture, almost all lysogens produced free phages in every experimental environment (except for prophage lambda carrying the ampicillin resistance gene; Figure 2). Addition of mitomycin C significantly increased the average amount of released phages (effect of mitomycin C: ampicillin treatment group: F_1,38_ = 61.44, p< 0.001; chloramphenicol treatment group: F_1,107_ = 25.84, p< 0.001; kanamycin treatment group: F_1,110_ = 81.39, p< 0.001). This indicates that, in the competition experiments above, addition of mitomycin C was probably associated with increased free phage production. Consistent with these treatments being associated with relatively high lysogen fitness in the competition experiment, we observed a positive association across all treatment groups between average lysogen fitness in the competition experiment and the concentration of free phages measured here (r=0.2, p=0.02; Figure S3). Note while free phage abundance was strongly influenced by mitomycin C, it also varied depending on which strains were present, being on average higher in cultures including ARG-carrying lysogens (Figure 2).

**Figure 2.**
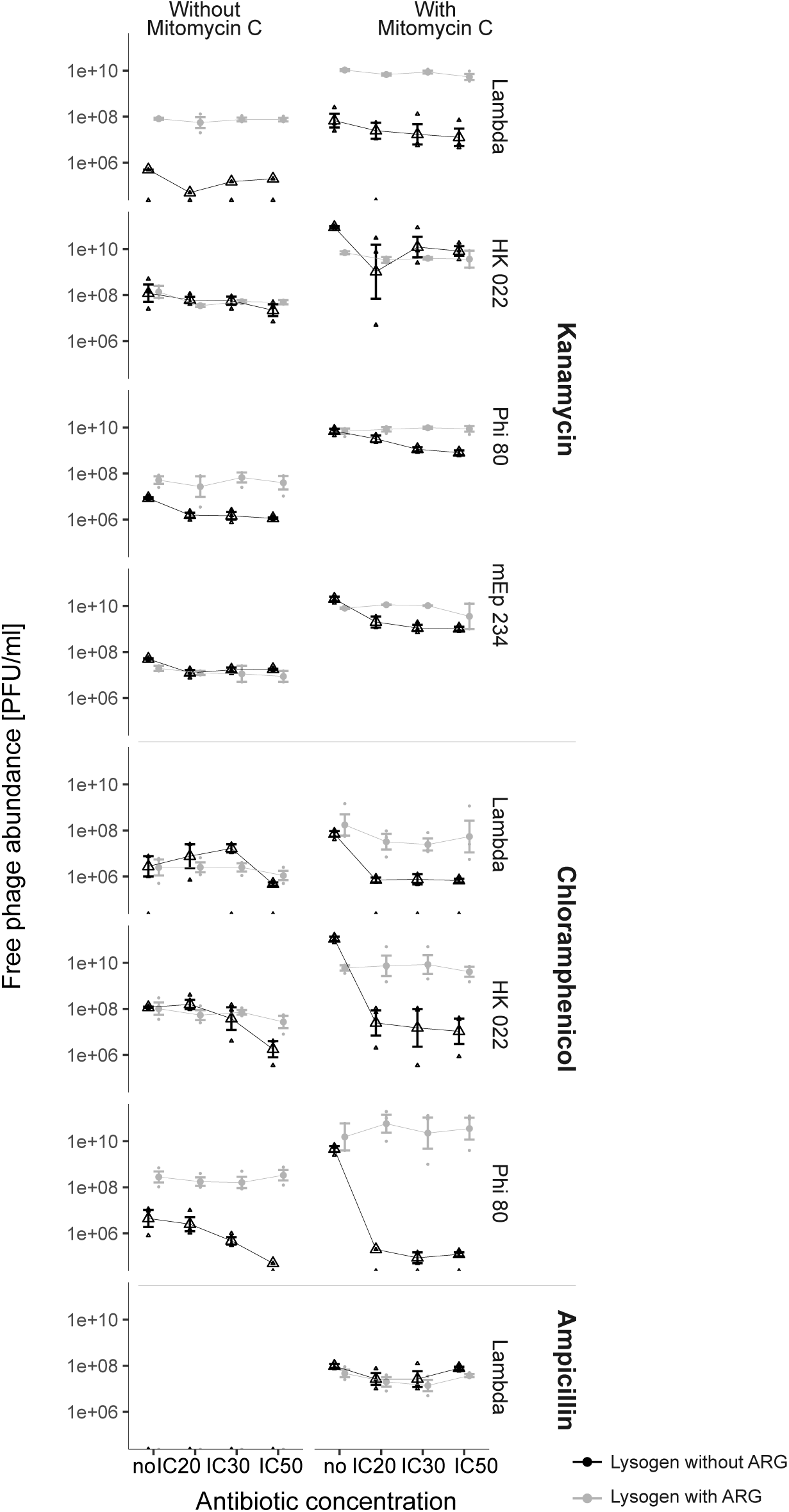
Free phage abundance in pure cultures of lysogens in the absence (left column) and presence (right column) of mitomycin C at different antibiotic concentrations (*x*-axis). Each row of panels gives data for ARG-free lysogens (black points) and ARG-carrying lysogens (grey points) for a different combination of antibiotic (given at far right) and prophage type (given at right). Shown are single data points as well as means ± s.e. from six replicate populations. Data-points for ampicillin/ without mitomycin C treatment are at *y*=0.

In the competition experiments, mitomycin C also caused a considerable reduction in total bacterial population densities (linear model - ampicillin treatment group: t=-6.3, p<0.001; chloramphenicol treatment group: t=-3.95, p<0.001; kanamycin treatment group: t=-2.03, p=0.04; Figure 3). This reduction in population density could have resulted from direct killing or growth inhibition by mitomycin C (for both the lysogen and the WT in each competition). Alternatively, it could have resulted from the effects of prophage induction and lysis (of prophage carrying cells upon switching to the lytic cycle, and of WT cells by released free phages). We found growth of prophage-free strains was not strongly reduced by mitomycin C at this concentration in pure culture (Figure S4). Prophage-carrying strains showed temporal growth dynamics upon exposure to mitomycin C in pure culture (Figure S4) consistent with prophage induction and abrupt lysis of a large fraction of the population, rather than constant growth inhibition by mitomycin C independent of the prophage. This suggests the reduced population densities in the presence of mitomycin C in competition cultures were probably caused by prophage induction, resulting in cell death of the induced lysogens and subsequent killing of the competing WT population by released phages. In support, using reduction in bacterial growth (RBG) assays, we confirmed the wild type was highly susceptible to growth inhibition by free phages (Supplementary material: Figure S5). Thus, experimental conditions supporting increased free phage production by lysogens were associated with increased competitive fitness of lysogens relative to the WT, even though both strains reached lower absolute abundances compared to when free phages were less abundant (Figure 3).

**Figure 3:**
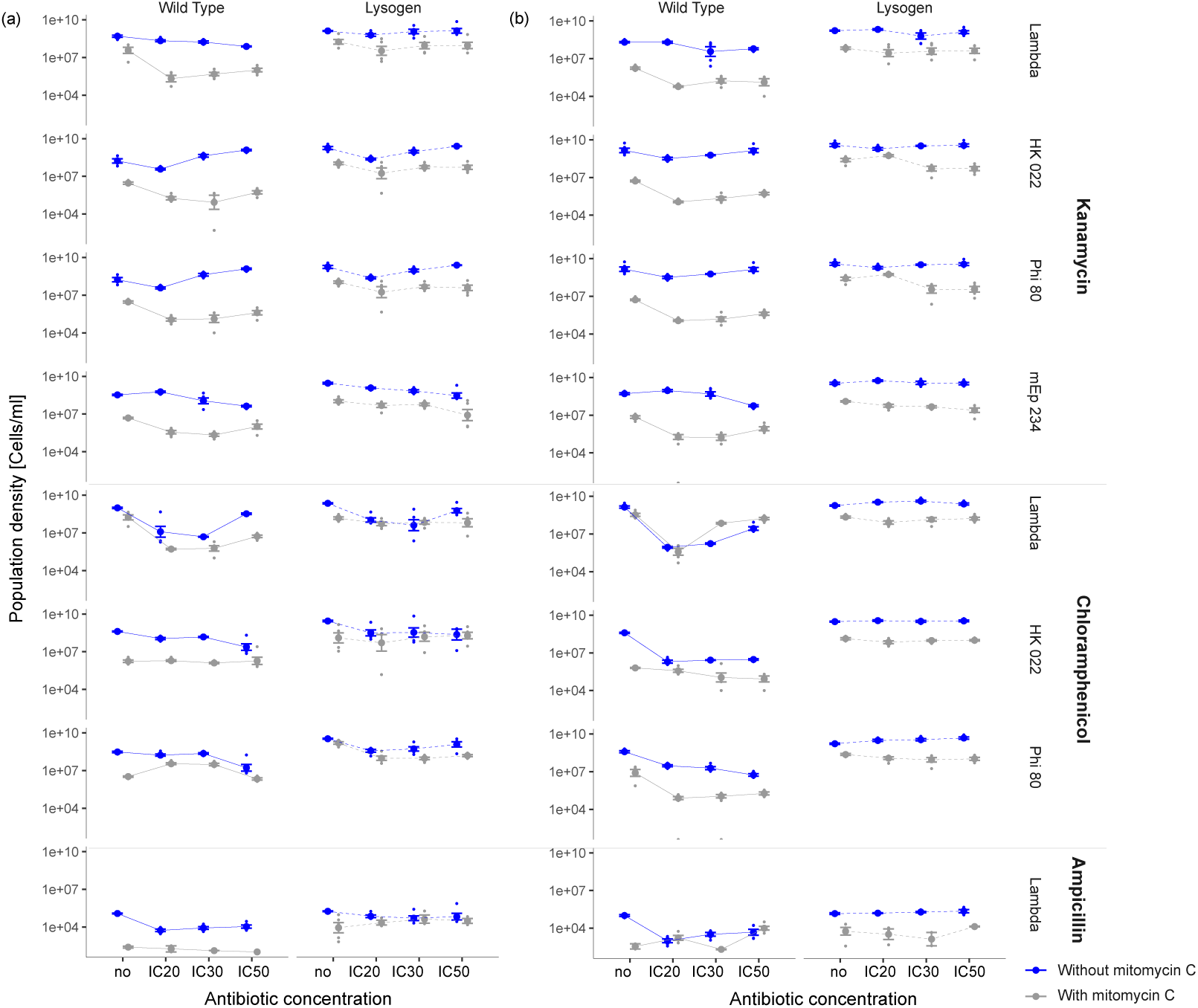
Bacterial population densities in competitions between (a) ARG-free lysogens and Wild Type bacteria and (b) ARG-carrying lysogens and Wild Type bacteria. Densities of Wild Type bacteria are shown in the left columns of (a) and (b), and densities of Lysogens are shown in the right columns of (a) and (b), at each antibiotic concentration (*x*-axis). Shown are single data points as well as means ± s.e. from six replicate populations. Blue/grey points are from competitions in the absence/presence of mitomycin C for different combinations of prophage type (given at the right of each row of panels) and antibiotic treatment (given at far right).

### The likelihood of ARG-transfer by prophages differs across environments

In 16 out of 320 pairwise competitions involving ARG-carrying lysogens, we detected putative WT lysogens that appeared to have acquired ARG-carrying prophages from the competing lysogen populations (Table 2). This was evidenced by their becoming antibiotic resistant, becoming inducible by mitomycin C (including production of free phages that could subsequently infect a susceptible WT clone), and their becoming resistant to a lytic version of the respective prophage (see Methods). 15 of these 16 WT lysogens emerged in the presence of antibiotics (antibiotic concentration effect in logistic regression: χ2_3_=8.52, p=0.04). Thirteen of the WT lysogens were detected in the presence of mitomycin C (mitomycin C effect in logistic regression: χ2_1_=8.36, p=0.004). The combination of intermediate antibiotic concentrations (IC20 and IC30) and the presence of mitomycin C were the same environments where lysogen fitness peaked in the competition experiments. These mostly fall into cluster one, representing environments where prophage carriage was more beneficial than ARG carriage (Figure 1c). Finally, most WT lysogens emerged in cultures with lambda prophages (12 out of 16), suggesting the probability of prophage-mediated ARG-transfer to a susceptible strain differs across prophage types (effect of prophage type in logistic regression: χ2_3_=12.73, p=0.005, Table 2).

**Table 2.** Treatment groups of all 16 populations where putative WT-lysogens (that appeared to have acquired a prophage during pairwise competition) were detected. Frequency: colony count for new WT lysogens in each culture as a percentage of the total colony count for the WT population.

| Phage genotype | Mitomycin C | Antibiotic | Concentration | Cluster | Frequency (%) |
| --- | --- | --- | --- | --- | --- |
| Lambda | Y | amp | IC50 | 1 | 0.06 |
| Lambda | Y | amp | IC30 | 1 | 0.06 |
| Lambda | Y | amp | IC30 | 1 | 0.08 |
| Lambda | Y | cm | IC30 | 1 | 0.06 |
| Lambda | Y | cm | IC30 | 1 | 0.16 |
| Lambda | Y | cm | IC30 | 1 | 0.05 |
| Lambda | Y | cm | IC20 | 1 | 0.09 |
| Lambda | Y | cm | IC20 | 1 | 0.08 |
| Lambda | Y | cm | IC20 | 1 | 0.04 |
| Lambda | Y | cm | IC50 | 2 | 0.37 |
| Lambda | Y | kan | IC50 | 1 | 0.04 |
| Lambda | Y | kan | IC30 | 1 | 0.04 |
| Lambda | Y | kan | IC30 | 1 | 6.67 |
| HK022 | N | kan | IC20 | 2 | 25 |
| Phi80 | Y | kan | IC30 | 1 | 15.38 |
| mEp234 | N | kan | 0 | 2 | 0.24 |
| HK022 | N | cm | IC30 | 3 | 0 |

### Prophage mediated transfer of resistance genes increases bacterial fitness in the presence of antibiotics

To determine how acquisition of ARG-carrying prophages in our main experiment affected bacterial fitness in these conditions, we carried out pairwise competitions between the new WT-lysogens (isolated from the competition experiment) and each of three different ancestral strains: (1) phage-free-ARG-free WT, (2) ARG-free lysogens with the same type of prophage, (3) ARG-carrying lysogens with the same type of prophage and ARG. We found fitness of new WT-lysogens relative to competing bacteria was affected by increasing antibiotic concentration, but this effect depended on the identity of the competing bacteria (competitor × environment interaction: ampicillin treatment group: F_2,50_ = 8.53, p<0.001; chloramphenicol treatment group: F_2,157_ = 134.99, p<0.001; kanamycin treatment group: F_2,184_ = 50.17, p<0.001; Figure 4). When they competed with either Wild-Type bacteria or ARG-free lysogens (Figure 4 left and middle columns), new WT lysogens had a fitness advantage in the presence but not the absence of antibiotics. By contrast, when they competed with ARG-carrying lysogens (Figure 4 right column), new WT lysogens had similar competitive fitness across the two environments. Thus, new WT lysogens behaved similarly to the bacteria they acquired prophages from, with a fitness advantage contingent on antibiotics and the presence of antibiotic-susceptible competitors.

**Figure 4.**
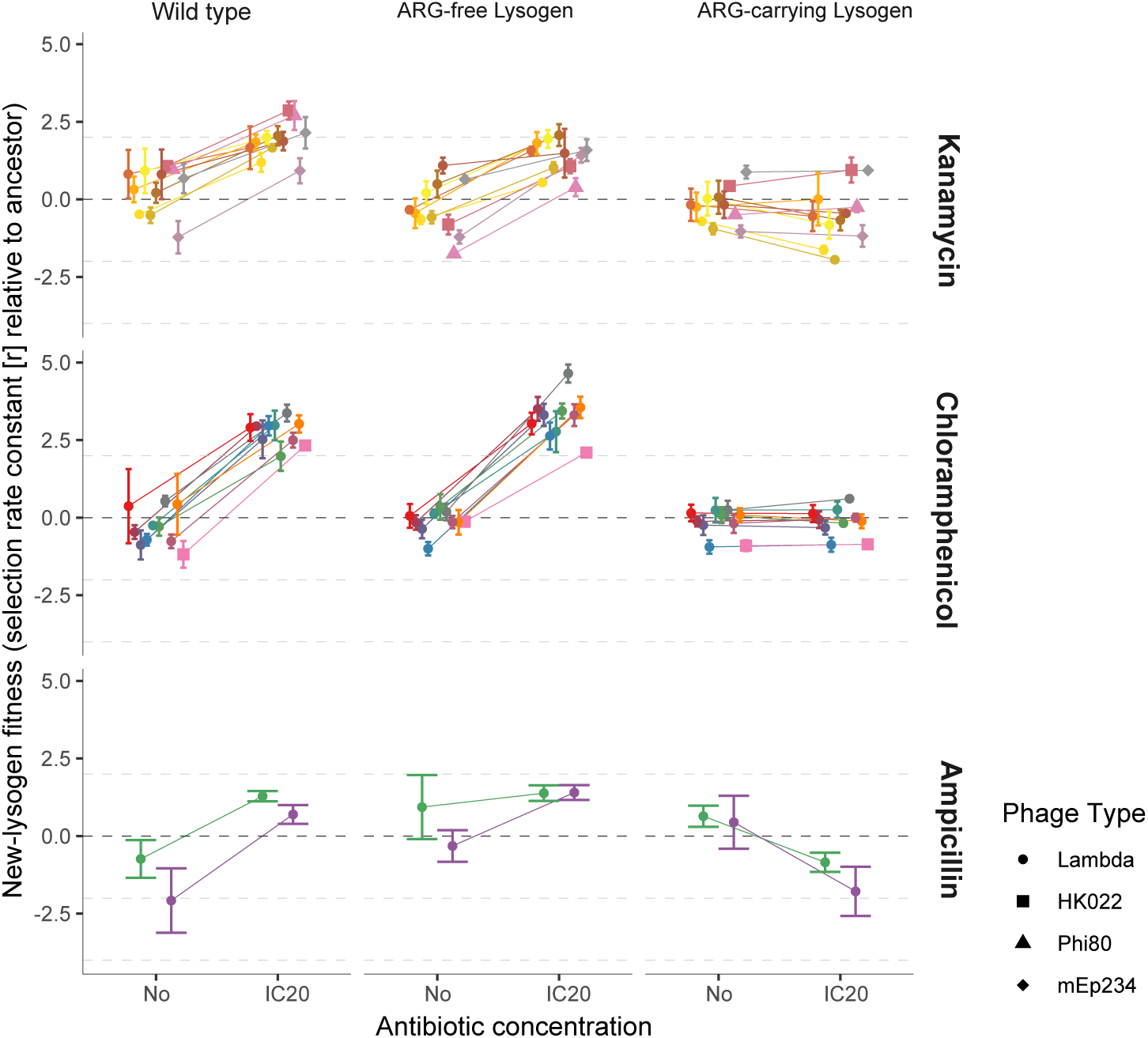
Competitive fitness of new WT-lysogens relative to each of three competing ancestral strains (left: phage-free, ARG-free Wild Type; middle: ARG-free lysogen with the same prophage type; right: ARG-carrying lysogen with the same prophage type), in the absence and presence of antibiotics (*x*-axis). Different symbols correspond to different prophage types (see legend). Points who means ± s.e. from three replicates. New WT lysogens have a higher fitness than the competing strain when *r* > 0. Each pair of connected points corresponds to a single new WT lysogen, isolated from one of the 16 populations where we detected new lysogens.

## DISCUSSION

We found fitness benefits resulting from prophage carriage peaked in different experimental environments compared to fitness advantages of prophage-encoded ARGs. Prophage-encoded ARGs, which were in most cases not associated with fitness costs in the absence of antibiotics (Andersson and Levin 1999), were more beneficial in environments where prophage induction was less frequent but antibiotics were present (cluster 3). In contrast, prophages were more beneficial than ARGs in environments where induction of the lytic cycle was more frequent (cluster 1). This beneficial effect of prophage carriage is likely driven by the release of free phages which, as determined in a separate experiment, was greatest in treatment groups where lysogen fitness was highest. Upon prophage induction, we expect the released phage particles infect and lyse competing, phage-susceptible WT cells (as observed in pure cultures - Figure S5; for a detailed review see (Harrison and Brockhurst 2017)). By selectively killing phage susceptible competitors, prophages can therefore increase the competitive fitness of their hosts (Bossi et al. 2003; De Paepe et al. 2016; Haaber et al. 2016). However, this benefit was (1) only visible at the population level, as individual cells are presumably killed upon induction of the lytic cycle (Figure 3) and (2) depended on being surrounded by phage-susceptible competitors, as lysogens did not grow better than phage-free strains in monoculture without antibiotics and mitomycin C (Figure S2). We observed formation of new WT lysogens in a small fraction of populations, indicating released phages can also drive genetic exchange. This observed horizontal transfer of prophage encoded ARGs was particularly favoured at sub-MIC conditions and provides new information about conditions in which prophages are likely to drive the transfer of resistance genes in bacterial communities.

The competitive advantage of lysogens over prophage-free competitors we observed supports the notion of a mutualistic, rather than parasitic, relationship between prophages and bacteria (Bondy-Denomy and Davidson 2014; Nanda et al. 2015; Fillol-Salom et al. 2019). In our experiments, the benefits of prophages appeared to result not only from their carriage of ARGs, but release of free phages that then lysed competing bacteria. Such killing of phage-susceptible competitors has been observed in other systems (Selva et al. 2009; Li et al. 2017). Release of free phages can also enable bacteria to acquire resistance genes from competing bacteria, as observed in our new WT lysogens. Thus, the prevalence of prophages in bacterial genomes may reflect a combination of direct competitive benefits in some environments and their role in exchange of genetic material (Jain et al. 1999; Ochman et al. 2000; Koonin et al. 2001). Prophages can constitute up to 20% of bacterial genomes and contribute significantly to sequence diversity among isolates (Casjens 2003). For instance, pathogenic *E. coli* serogroup O157 contain between 15 and 18 different prophages, which account for most of the increased genome size (5.5 Mb) compared to K-12 MG1655 (4.6 Mb, (Perna et al. 2001)).

Our key finding that prophages were beneficial across different environments applied across multiple phage types and antibiotics. Despite this, we observed variation of lysogen fitness across phage types in our chloramphenicol treatment, and a cost of prophage carriage for lambda lysogens in the presence of ampicillin and mitomycin C. This suggests lysogen fitness is also influenced by an interaction between the type of prophage and the type of antibiotic. There are multiple possible mechanisms by which such an interaction could arise, and an interesting avenue for future work would be to investigate this for a much larger range of prophage types and antibiotics, including multiple antibiotics from each class. We nevertheless speculate that the relatively high fitness of prophage-free bacteria in the presence of ampicillin may reflect the mechanism of ampicillin resistance. For β-lactams such as ampicillin, resistance is often mediated via β-lactamases that deactivate the antibiotic, reducing the effective concentration and potentially diminishing inhibitory effects on susceptible cells in mixed cultures (Medaney et al. 2016).

Some key properties of our experimental conditions are likely to also apply in nature. Many pathogenic *E. coli* harbour lambdoid prophages (Casjens 2003), and co-occurrence of closely related enterobacteria that differ in their prophage repertoire are common in gastrointestinal tracts. About 25% of 243 tested coliphages are able to transduce ARGs (encoding ampicillin, chloramphenicol, kanamycin or tetracycline resistance) to the laboratory strain *E. coli* ATCC 13706 (Shousha et al. 2015). Moreover, in animal farms, sub-MIC concentrations of antibiotics are likely common as a result of antibiotic application as growth promoters and prophylactics (Van Boeckel et al. 2015). Thus, we can expect conditions that would favour bacterial acquisition of such ARG-carrying prophages to be widespread. Understanding the drivers of lysogen fitness, such as variable free phage production depending on abiotic conditions as we observed, is therefore important for predicting the risk that prophage-encoded ARGs could spread from agricultural or natural environments to human microbiota or pathogens.

Another key implication of our results is that lysogenization of phage-susceptible cells by ARG-carrying prophages appears to be most likely in the presence of antibiotics and in conditions where free phage abundance is high. There are at least two possible explanations. First, the relatively high benefits of acquiring prophages and ARGs in these conditions presumably increased their final abundance and the probability of us detecting them. For ARGs, we observed stronger fitness benefits in the presence of antibiotics, both in our main competition assay and in experiments with new WT lysogens. Independently of ARGs, the benefits of prophage acquisition were higher in the presence of mitomycin C and antibiotics. This may result from the relatively high abundances of free phages in these treatment groups, indicated by our phage-production experiment. In such conditions, we can expect the benefits of prophage acquisition stemming from immunity to super-infection (Lwoff 1953) to be relatively high. Consistent with this, previous work showed lysogeny of initially phage-susceptible bacteria can increase their competitive fitness relative to initially-prophage-carrying bacteria (Gama et al. 2013). Note that, as well as mitomycin C, other antibiotics or drugs can induce the lytic cycle and trigger the release of free phages. This has been observed for prophages of *Staphylococcus aureus* (Ubeda et al. 2005; Maiques et al. 2006) and the virus-like gene transfer agent VSH-1 in the swine pathogen *Brachyspira hyodysenteriae* (Matson et al. 2005; Stanton et al. 2008). We therefore speculate that exposure to antibacterial compounds will often promote new lysogens, either via direct selection for prophage-encoded resistance genes or indirectly by inducing free phage production, which in turn selects for prophage-derived resistance to superinfection.

A second, non-exclusive explanation for the bias of new lysogen formation toward antibiotic-and mitomycin C treatments is that the frequency of lysogenization events per phage-bacteria contact may be higher in these treatment groups. For instance, in conditions that are sub-optimal for bacteria, such as low temperature (Obuchowski et al. 1997) or starvation, phage lambda is biased toward the lysogenic over the lytic cycle (Herskowitz and Hagen 1980). Similarly, lysogeny is favoured when the multiplicity of infection (MOI) is high (Kourilsky 1973; Kobiler et al. 2005). Our finding that free phage production was higher in the presence of antibiotics and mitomycin C (presumably due to more frequent induction of the lytic cycle) does not exclude the possibility that formation of new lysogens per cell contact was also more frequent in these conditions.

### Conclusion

We provide empirical evidence that the benefits to lysogens of carrying prophages and prophage-encoded ARGs vary independently across abiotic conditions, with the release of free phages from lysogens entering the lytic cycle appearing to be a key driver of prophage-associated fitness benefits. By disentangling the benefits of prophages and ARGs, this expands our understanding of selection on prophages in the presence of phage-susceptible competitors (Gama et al. 2013; Davies et al. 2016). This study provides new information about the evolutionary biology of mobile genetic elements by showing that selection on MGEs and beneficial genes encoded thereon varies independently across environments. In addition, we show that sub-MIC concentrations of antibiotics favour the horizontal transfer of phage-encoded resistance genes to phage- and antibiotic susceptible strains. This improves our understanding of the conditions in which prophages are likely to drive transfer of ARGs in bacterial communities.

## Acknowledgements

We thank Iris Wurmitzer for help in the laboratory and Tobias Bergmiller for support on recombineering. This project has received funding from the European Union’s Horizon 2020 research and innovation programme under the Marie Sklodowska-Curie grant agreement No 794447-ProphARG given to CCW. Construction of phages and lysogens was carried out under the project ‘Genetic Diversity of Hosts and Parasites’ (grant no. 0-21248-08), funded by the Competence Center Environment and Sustainability of the ETH, given to Sebastian Bonhoeffer.

## References

Andersson, D. I. and D. Hughes. 2014. Microbiological effects of sublethal levels of antibiotics. Nat Rev Microbiol 12:465–478.

Andersson, D. I. and B. R. Levin. 1999. The biological cost of antibiotic resistance. Curr Opin Microbiol 2:489–493.

Balcazar, J. L. 2014. Bacteriophages as Vehicles for Antibiotic Resistance Genes in the Environment. Plos Pathogens 10.

Bondy-Denomy, J. and A. R. Davidson. 2014. When a Virus is not a Parasite: The Beneficial Effects of Prophages on Bacterial Fitness. J Microbiol 52:235–242.

Bossi, L., J. A. Fuentes, G. Mora, and N. Figueroa-Bossi. 2003. Prophage contribution to bacterial population dynamics. J Bacteriol 185:6467–6471.

Brenciani, A., A. Bacciaglia, C. Vignaroli, A. Pugnaloni, P. E. Varaldo, and E. Giovanetti. 2010. Phi m46.1, the Main Streptococcus pyogenes Element Carrying mef(A) and tet(O) Genes. Antimicrob Agents Ch 54:221–229.

Calero-Caceres, W., M. Ye, and J. L. Balcazar. 2019. Bacteriophages as Environmental Reservoirs of Antibiotic Resistance. Trends Microbiol 27:570–577.

Carlson, K. 2005. Working with bacteriophages: common techniques and methodological approaches. Pp. 429-484 *in* E. Kutter, and A. Sulakvelidze, eds. Bacteriophages: Biology and applications. GRC Press, Boca Raton, Florida.

Casjens, S. 2003. Prophages and bacterial genomics: what have we learned so far? Mol Microbiol 49:277–300.

Datsenko, K. A. and B. L. Wanner. 2000. One-step inactivation of chromosomal genes in Escherichia coli K-12 using PCR products. P Natl Acad Sci USA 97:6640–6645.

Davies, E. V., C. E. James, I. Kukavica-Ibrulj, R. C. Levesque, M. A. Brockhurst, and C. Winstanley. 2016. Temperate phages enhance pathogen fitness in chronic lung infection. Isme J 10:2553–2555.

De Paepe, M., L. Tournier, E. Moncaut, O. Son, P. Langella, and M. A. Petit. 2016. Carriage of. Latent Virus Is Costly for Its Bacterial Host due to Frequent Reactivation in Monoxenic Mouse Intestine. Plos Genetics 12.

Debroas, D. and C. Siguret. 2019. Viruses as key reservoirs of antibiotic resistance genes in the environment. The ISME Journal.

Fillol-Salom, A., A. Alsaadi, J. A. M. de Sousa, L. Zhong, K. R. Foster, E. P. C. Rocha, J. R. Penades, H. Ingmer, and J. Haaber. 2019. Bacteriophages benefit from generalized transduction. Plos Pathogens 15.

Gama, J. A., A. M. Reis, I. Domingues, H. Mendes-Soares, A. M. Matos, and F. Dionisio. 2013. Temperate Bacterial Viruses as Double-Edged Swords in Bacterial Warfare. Plos One 8.

Gullberg, E., L. M. Albrecht, C. Karlsson, L. Sandegren, and D. I. Andersson. 2014. Selection of a multidrug resistance plasmid by sublethal levels of antibiotics and heavy metals. Mbio 5:e01918–01914.

Haaber, J., J. J. Leisner, M. T. Cohn, A. Catalan-Moreno, J. B. Nielsen, H. Westh, J. R. Penades, and H. Ingmer. 2016. Bacterial viruses enable their host to acquire antibiotic resistance genes from neighbouring cells. Nat Commun 7.

Harrison, E. and M. A. Brockhurst. 2017. Ecological and Evolutionary Benefits of Temperate Phage: What Does or Doesn’t Kill You Makes You Stronger. BioEssays : news and reviews in molecular, cellular and developmental biology.

Herskowitz, I. and D. Hagen. 1980. The lysis-lysogeny decision of phage lambda: explicit programming and responsiveness. Annual review of genetics 14:399–445.

Jain, R., M. C. Rivera, and J. A. Lake. 1999. Horizontal gene transfer among genomes: The complexity hypothesis. P Natl Acad Sci USA 96:3801–3806.

Kobiler, O., A. Rokney, N. Friedman, D. L. Court, J. Stavans, and A. B. Oppenheim. 2005. Quantitative kinetic analysis of the bacteriophage lambda genetic network. Proc Natl Acad Sci U S A 102:4470–4475.

Koonin, E. V., K. S. Makarova, and L. Aravind. 2001. Horizontal gene transfer in prokaryotes: quantification and classification. Annu Rev Microbiol 55:709–742.

Kourilsky, P. 1973. Lysogenization by bacteriophage lambda. I. Multiple infection and the lysogenic response. Mol Gen Genet 122:183–195.

Lenski, R. E., M. R. Rose, S. C. Simpson, and S. C. Tadler. 1991. Long-Term Experimental Evolution in Escherichia-Coli .1. Adaptation and Divergence during 2,000 Generations. Am Nat 138:1315–1341.

Li, X. Y., T. Lachnit, S. Fraune, T. C. G. Bosch, A. Traulsen, and M. Sieber. 2017. Temperate phages as self-replicating weapons in bacterial competition. J R Soc Interface 14.

Lwoff, A. 1953. Lysogeny. Bacteriol Rev 17:269–337.

Maiques, E., C. Ubeda, S. Campoy, N. Salvador, I. Lasa, R. P. Novick, J. Barbe, and J. R. Penades. 2006. beta-lactam antibiotics induce the SOS response and horizontal transfer of virulence factors in Staphylococcus aureus. J Bacteriol 188:2726–2729.

Mann, N. H. 2005. The third age of phage. PLoS Biol 3:e182.

Matson, E. G., M. G. Thompson, S. B. Humphrey, R. L. Zuerner, and T. B. Stanton. 2005. Identification of genes of VSH-1, a prophage-like gene transfer agent of Brachyspira hyodysentetiae. J Bacteriol 187:5885–5892.

Medaney, F., T. Dimitriu, R. J. Ellis, and B. Raymond. 2016. Live to cheat another day: bacterial dormancy facilitates the social exploitation of beta-lactamases. Isme J 10:778–787.

Modi, S. R., H. H. Lee, C. S. Spina, and J. J. Collins. 2013. Antibiotic treatment expands the resistance reservoir and ecological network of the phage metagenome. Nature 499:219-+.

Nanda, A. M., K. Thormann, and J. Frunzke. 2015. Impact of Spontaneous Prophage Induction on the Fitness of Bacterial Populations and Host-Microbe Interactions. J Bacteriol 197:410–419.

Obuchowski, M., Y. Shotland, S. Koby, H. Giladi, M. Gabig, G. Wegrzyn, and A. B. Oppenheim. 1997. Stability of CII is a key element in the cold stress response of bacteriophage lambda infection. J Bacteriol 179:5987–5991.

Ochman, H., J. G. Lawrence, and E. A. Groisman. 2000. Lateral gene transfer and the nature of bacterial innovation. Nature 405:299–304.

Otsuji, N., M. Sekiguchi, T. Iijima, and Y. Takagi. 1959. Induction of Phage Formation in the Lysogenic Escherichia-Coli K-12 by Mitomycin-C. Nature 184:1079–1080.

Palmieri, C., M. S. Princivalli, A. Brenciani, P. E. Varaldo, and B. Facinelli. 2011. Different genetic elements carrying the tet(W) gene in two human clinical isolates of Streptococcus suis. Antimicrob Agents Chemother 55:631–636.

Paul, J. H. 2008. Prophages in marine bacteria: dangerous molecular time bombs or the key to survival in the seas? Isme J 2:579–589.

Perna, N. T., G. Plunkett, V. Burland, B. Mau, J. D. Glasner, D. J. Rose, G. F. Mayhew, P. S. Evans, J. Gregor, H. A. Kirkpatrick, G. Posfai, J. Hackett, S. Klink, A. Boutin, Y. Shao, L. Miller, E. J. Grotbeck, N. W. Davis, A. Limk, E. T. Dimalanta, K. D. Potamousis, J. Apodaca, T. S. Anantharaman, J. Y. Lin, G. Yen, D. C. Schwartz, R. A. Welch, and F. R. Blattner. 2001. Genome sequence of enterohaemorrhagic Escherichia coli O157 : H7. Nature 409:529–533.

Poullain, V., S. Gandon, M. A. Brockhurst, A. Buckling, and M. E. Hochberg. 2008. The evolution of specificity in evolving and coevolving antagonistic interactions between a bacteria and its phage. Evolution 62:1–11.

Pricer, W. E., Jr. and A. Weissbach. 1964. The Effect of Lysogenic Induction with Mitomycin C on the Deoxyribonucleic Acid Polymerase of Escherichia Coli K12-Lambda. J Biol Chem 239:2607–2612.

Qi, Q., M. Toll-Riera, K. Heilbron, G. M. Preston, and R. C. MacLean. 2016. The genomic basis of adaptation to the fitness cost of rifampicin resistance in Pseudomonas aeruginosa. Proceedings. Biological sciences 283.

Refardt, D. 2011. Within-host competition determines reproductive success of temperate bacteriophages. Isme J 5:1451–1460.

Regoes, R. R., C. Wiuff, R. M. Zappala, K. N. Garner, F. Baquero, and B. R. Levin. 2004. Pharmacodynamic functions: a multiparameter approach to the design of antibiotic treatment regimens. Antimicrob Agents Chemother 48:3670–3676.

Rohwer, F. 2003. Global phage diversity. Cell 113:141.

Selva, L., D. Viana, G. Regev-Yochay, K. Trzcinski, J. M. Corpa, I. Lasa, R. P. Novick, and J. R. Penades. 2009. Killing niche competitors by remote-control bacteriophage induction. P Natl Acad Sci USA 106:1234–1238.

Shousha, A., N. Awaiwanont, D. Sofka, F. J. M. Smulders, P. Paulsen, M. P. Szostak, T. Humphrey, and F. Hilbert. 2015. Bacteriophages Isolated from Chicken Meat and the Horizontal Transfer of Antimicrobial Resistance Genes. Appl Environ Microb 81:4600–4606.

Sprouffske, K. and A. Wagner. 2016. Growthcurver: an R package for obtaining interpretable metrics from microbial growth curves. Bmc Bioinformatics 17:172.

Stanton, T. B., S. B. Humphrey, V. K. Sharma, and R. L. Zuerner. 2008. Collateral effects of antibiotics: Carbadox and metronidazole induce VSH-1 and facilitate gene transfer among Brachyspira hyodysenterae strains. Appl Environ Microb 74:2950–2956.

Ubeda, C., E. Maiques, E. Knecht, I. Lasa, R. P. Novick, and J. R. Penades. 2005. Antibiotic-induced SOS response promotes horizontal dissemination of pathogenicity island-encoded virulence factors in staphylococci. Mol Microbiol 56:836–844.

Van Boeckel, T. P., C. Brower, M. Gilbert, B. T. Grenfell, S. A. Levin, T. P. Robinson, A. Teillant, and R. Laxminarayan. 2015. Global trends in antimicrobial use in food animals. Proc Natl Acad Sci U S A 112:5649–5654.

