## Supplementary material for "FITNESS BENEFITS TO BACTERIA OF CARRYING PROPHAGES AND PROPHAGE-ENCODED ANTIBIOTIC-RESISTANCE GENES PEAK IN DIFFERENT ENVIRONMENTS"

**Table S1** Primer sequences that were used to amplify resistance cassettes and exchange them with non-essential genes in prophages.

| Amplified resistance cassette | Homology extension | Primer sequence (bold: homology extension) |
|---|---|---|
| *neo* | lambda *bor* | 5’-**ACGATTCTGCGAACTTCAAAAAGCATCGGGAATAACACC**TCACGCTGCCGCAAGCACTC-3’<br>5’-**TGCAGATAGAGTTGCCCATATCGATGGGCAACTCATGCAA**AGCGCTTTTGAAGCTGGGGTG-3’ |
| *neo* | HK022 *cor* | 5’-**GTATCTTCGAAGCTACTGAATACCTCTGGATACTATCAAA**TCACGCTGCCGCAAGCACTC-3’<br>5’-**AAAACCCCGCCGAGGCGAGGTTTGCATTGAAAATCATGAG**AGCGCTTTTGAAGCTGGGGTG-3’ |
| *neo* | Phi80 *cor* | 5’-**TTTCTCCAGGCAATAAAAAAACCCCGCCGGAGCGAGGTTT**TCACGCTGCCGCAAGCACTC-3’<br>5’-**CCAATTATACCCACCTCTTTCATGTTGGTATTGTCTAAGT**AGCGCTTTTGAAGCTGGGGTG-3’ |
| *neo* | mEp234 *cor* | 5’-**GTATCTTCGAAGCTACTGAATACCTCTGGATACTATCAAA**TCACGCTGCCGCAAGCACTC-3’<br>5’-**AAAACCCCGCCGAGGCGAGGTTTGCATTGAAAATCATGAG**AGCGCTTTTGAAGCTGGGGTG-3’ |
| *cat* | lambda *bor* | 5’-**TACGATTCTGCGAACTTCAAAAAGCATCGGGAATAACACC**CGCGCCTACCTGTGACGGAA-3’<br>5’-**TGCAGATAGAGTTGCCCATATCGATGGGCAACTCATGCAA**AAATGGCGCGCCTTACGCCC-3’ |
| *cat* | HK022 *cor* | 5’-**GCTGTCTGGTTGCGCTGGTGTACTTGAGAAACAGAAACCG**CGCGCCTACCTGTGACGGAA-3’<br>5’-**TCCGGCCCGGTATTGCGTCTGGTTGTTTTGCTTTCGAACG**AAATGGCGCGCCTTACGCCC-3’ |
| *cat* | Phi80 *cor* | 5’-**TTTCTCCAGGCAATAAAAAAACCCCGCCGGAGCGAGGTTT**CGCGCCTACCTGTGACGGAA-3’<br>5’-**CCAATTATACCCACCTCTTTCATGTTGGTATTGTCTAAGT**AAATGGCGCGCCTTACGCCC-3’ |
| *bla* | lambda *bor* | 5’-**TACGATTCTGCGAACTTCAAAAAGCATCGGGAATAACACC**CATTCAAATATGTATCCGCTC-3’<br>5’-**TGCAGATAGAGTTGCCCATATCGATGGGCAACTCATGCAA**AGAGTTGGTAGCTCTTGATC-3’ |

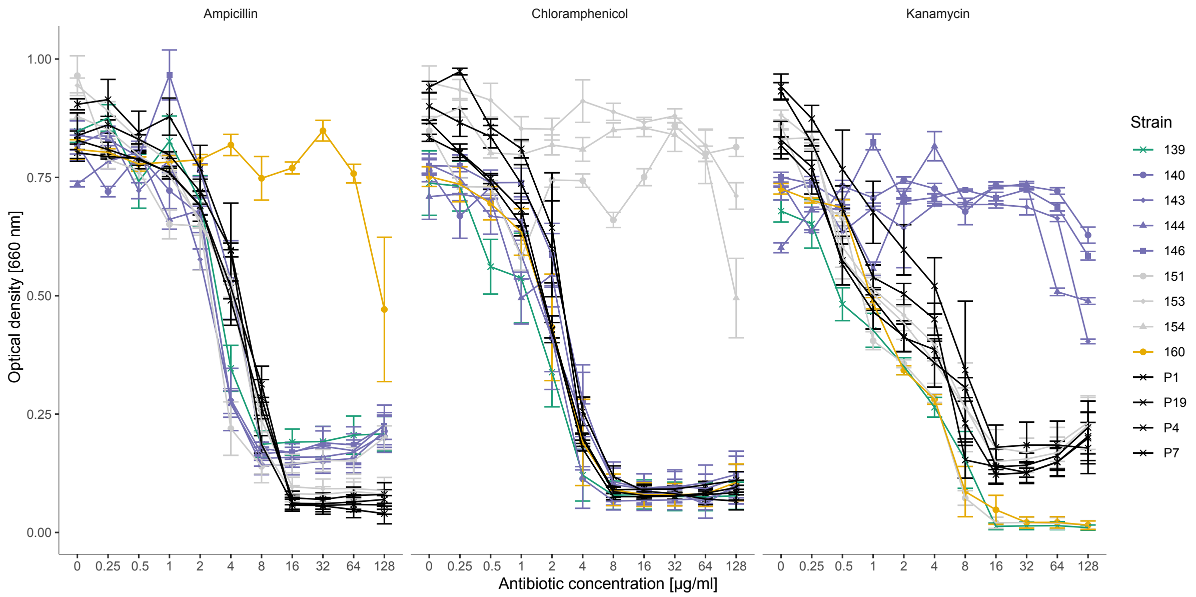


**Figure S1** Bacterial growth measured at multiple concentrations of three antibiotics (given in panel labels at top). We measured bacterial growth as optical density at 660 nm after 24h of incubation. Strains (see legend; described in more detail in Table 1 of the main text) are coded according to resistance gene and phage type: green: WT (no ARG, no prophage), purple: kanamycin, grey: chloramphenicol, orange: ampicillin, black: no ARG. Prophage type is indicated by the symbol shape - circle: lambda+, diamond: HK022, triangle: Phi80, square: mep234, cross: no prophage (see legend). Shown are means ± s.e. from six replicate populations.

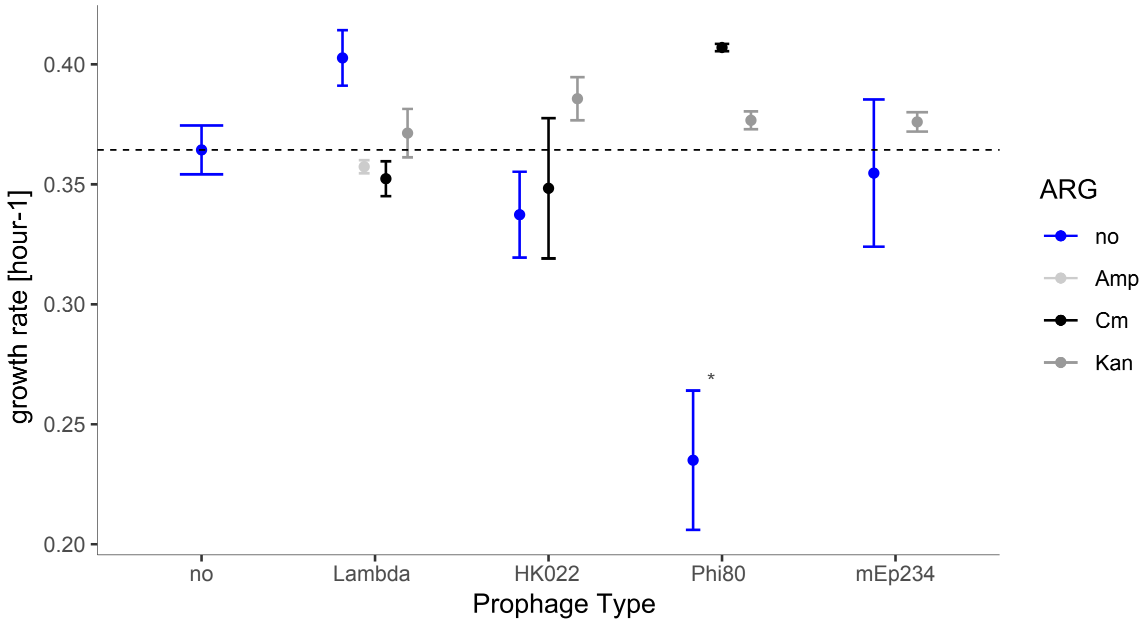


**Figure S2** Growth rate [r] of each lysogen in the absence of antibiotics and mitomycin C, arranged by prophage type (*x*-axis) and antibiotic resistance gene (see legend). Growth rate [r] was extracted from 24-hour growth curves. The asterisk indicates the one lysogen (strain P7) which grew significantly slower than the WT (Welch's *t*-test: $t_{3.5}$=-4.55, p=0.0005). Growth rates of all other lysogens were not significantly different from the WT (p>α in pairwise *t*-tests after sequential Bonferroni correction). Points show means ± s.e.; the dashed line shows the mean growth of WT.

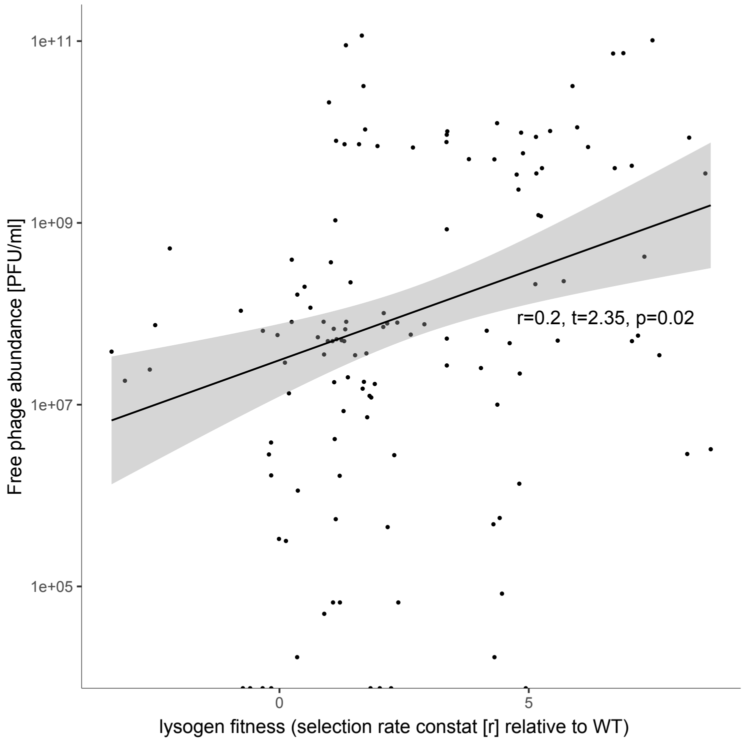


**Figure S3** Correlation between log-transformed free phage abundance (measured after four hours in pure cultures of lysogens; *y*-axis) and lysogen relative fitness (measured in competition with the prophage-free WT; *x*-axis). Each point represents the mean for a different treatment group (combination of lysogen, prophage type, presence/absence of mitomycin C, antibiotic and antibiotic concentration).

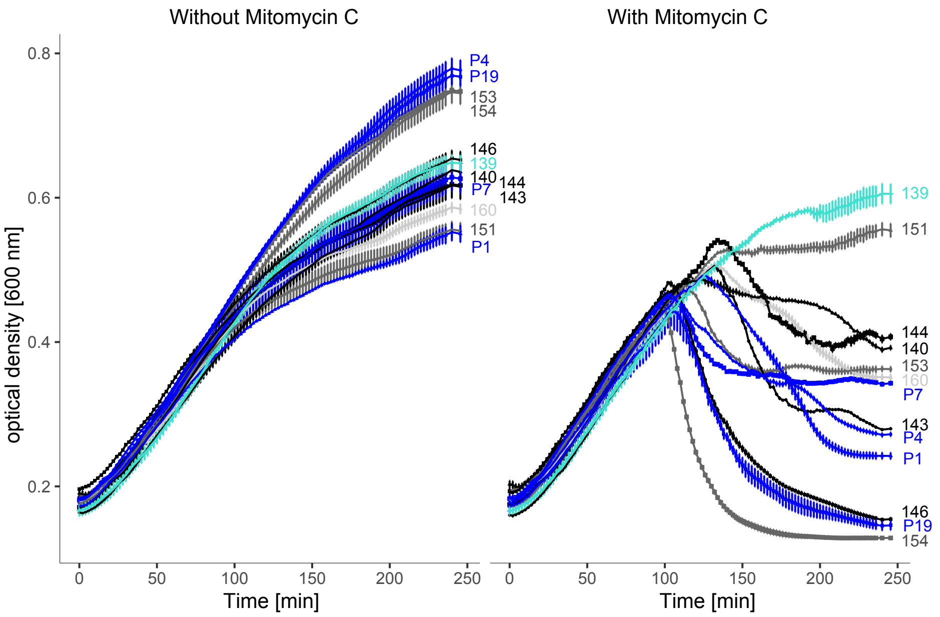


**Figure S4** Population density of each strain over time (20 h) in the presence of mitomycin C (0.5 $\mu$g/ml). Strains are labelled at right (139 is the WT, blue lines are lysogens without ARGs, light grey lines are lysogens with ampicillin resistance, dark grey lines are lysogens with chloramphenicol resistance, black lines are lysogens with kanamycin resistance; see Table 1 in the main text for more details).

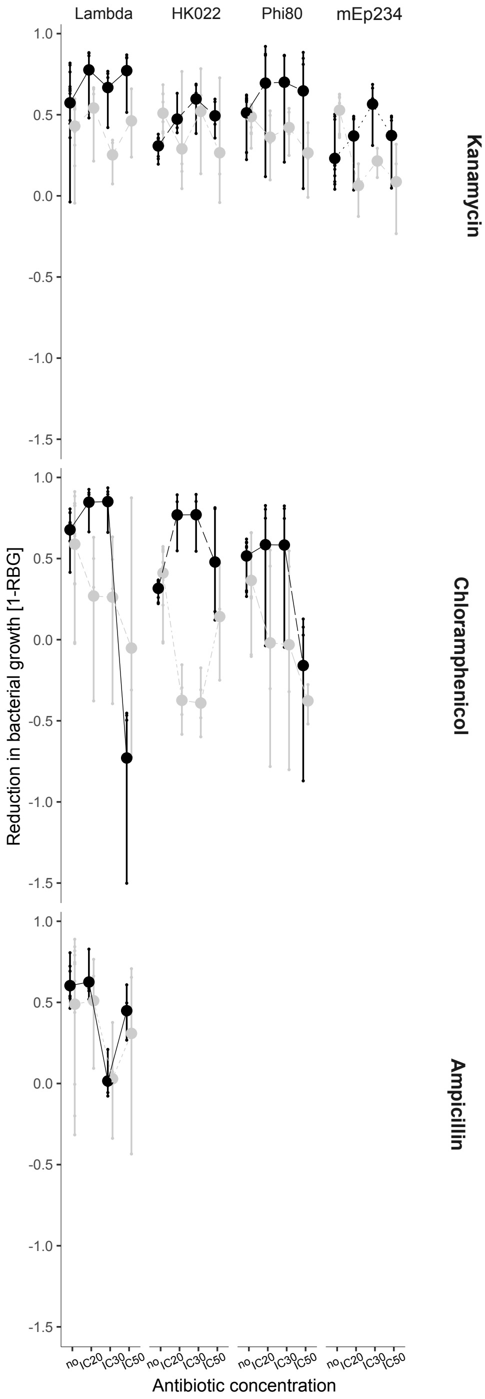


**Figure S5** Reduction in bacterial growth (1-RBG) imposed by free phages for each phage genotype (columns, labelled at top) for each antibiotic (rows, labelled at right), without an ARG = black, with ARG = grey at four antibiotic concentrations (*x*-axis). Higher RBG-values indicate stronger inhibition of bacterial growth imposed by the phage.
